# Crosslinking actin networks produces compressive force

**DOI:** 10.1101/614453

**Authors:** Rui Ma, Julien Berro

## Abstract

Actin has been shown to be essential for clathrin-mediated endocytosis in yeast. However, actin polymerization alone is likely insufficient to produce enough force to deform the membrane against the huge turgor pressure of yeast cells. In this paper, we used Brownian dynamics simulations to demonstrate that crosslinking of a meshwork of non-polymerizing actin filaments is able to produce compressive forces. We show that the force can be up to thousands of piconewtons if the crosslinker has a high stiffness. The force decays over time as a result of crosslinker turnover, and is a result of converting chemical binding energy into elastic energy.

## Introduction

Polymerization of actin filaments provides the direct driving force for many cellular processes, such as the movement of cells and bacteria (Mitchison and Cramer 1996; Pollard and Cooper 2009; Loisel et al. 1999). Actin filaments can also work with myosin motors to power processes like muscle contraction (Rayment et al. 1993; Spudich and Watt 1971) and cytokinesis (Pollard 2010; Pollard and Wu 2010). During clathrin-mediated endocytosis (CME), a flat patch of membrane is deformed towards the cytoplasm to form a vesicle. In yeast cells, this process is opposed by a high turgor pressure of 1-1.5 MPa (Minc, Boudaoud, and Chang 2009; Basu, Munteanu, and Chang 2014; Atilgan et al. 2015) and actin network is necessary for successful CME (Kaksonen, Toret, and Drubin 2006; Mooren, Galletta, and Cooper 2012). However, actin polymerization-based and motor-mediated force production are unable to meet the force budget required to internalize a vesicle, which is up to ∼3000 pN due to the high turgor pressure (Dmitrieff and Nédélec 2015; Lacy et al. 2018). Assuming each actin filament can generate a force of 1pN at their polymerizing end (Footer et al. 2007), the maximum force provided by polymerization alone is less than ∼200pN given the number of actin filaments involved in CME is no more than 200 (Berro, Sirotkin, and Pollard 2010; Sirotkin et al. 2010). The detachment rate of myosin-I motors involved in CME significantly increases once the loading force is beyond 2pN (Laakso et al. 2008), suggesting a maximum force of ∼600pN can be produced by motor-filament interaction given the number of myosin-I motors is ∼300 (Sirotkin et al. 2010). Therefore, other force production mechanisms complementary to polymerization and motor-filament interaction need to be investigated.

Actin crosslinking protein fimbrin is the second most abundant protein involved in CME next to actin (Sirotkin et al. 2010). Though fimbrin is a passive linker that does not convert the chemical energy of ATP hydrolysis into mechanical work like molecular motors do, we have shown that its binding chemical energy can be converted into elastic energy upon crosslinking actin filaments (Ma and Berro 2018). We wondered whether crosslinking may also be able to directly produce force in certain conditions. We hypothesized that an object in the middle of an crosslinked actin meshwork would be subject to compressive forces. Providing endocytosis is fast, this mechanism could be a non-negligible source of force production, even if it is transient.

In this paper, we study the force produced by crosslinking a network of non-polymerizing actin filaments around an elastic cylinder. We find that the force is compressive and decays over time due to crosslinker turnover. The magnitude of the force can be up to thousands of pico-newton if the stiffness of the crosslinker is high.

## Model

We place a total number of *N*_fil_ = 140 actin filaments around a purely elastic cylinder of radius *R* and stiffness *κ*_cyl_. If a filament penetrates into the cylinder, such that the filament’s nearest distance to the cylinder’s central axis *D* < *R*, it experiences a repulsive force **F** = *F* **d** with a magnitude *F* that is proportional to the penetration depth

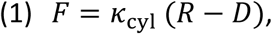

and **d** is the unit vector pointing outward along the axis that defines the nearest distance between the filament and the central axis of the cylinder (Fig.1 a).

**Figure 1:**
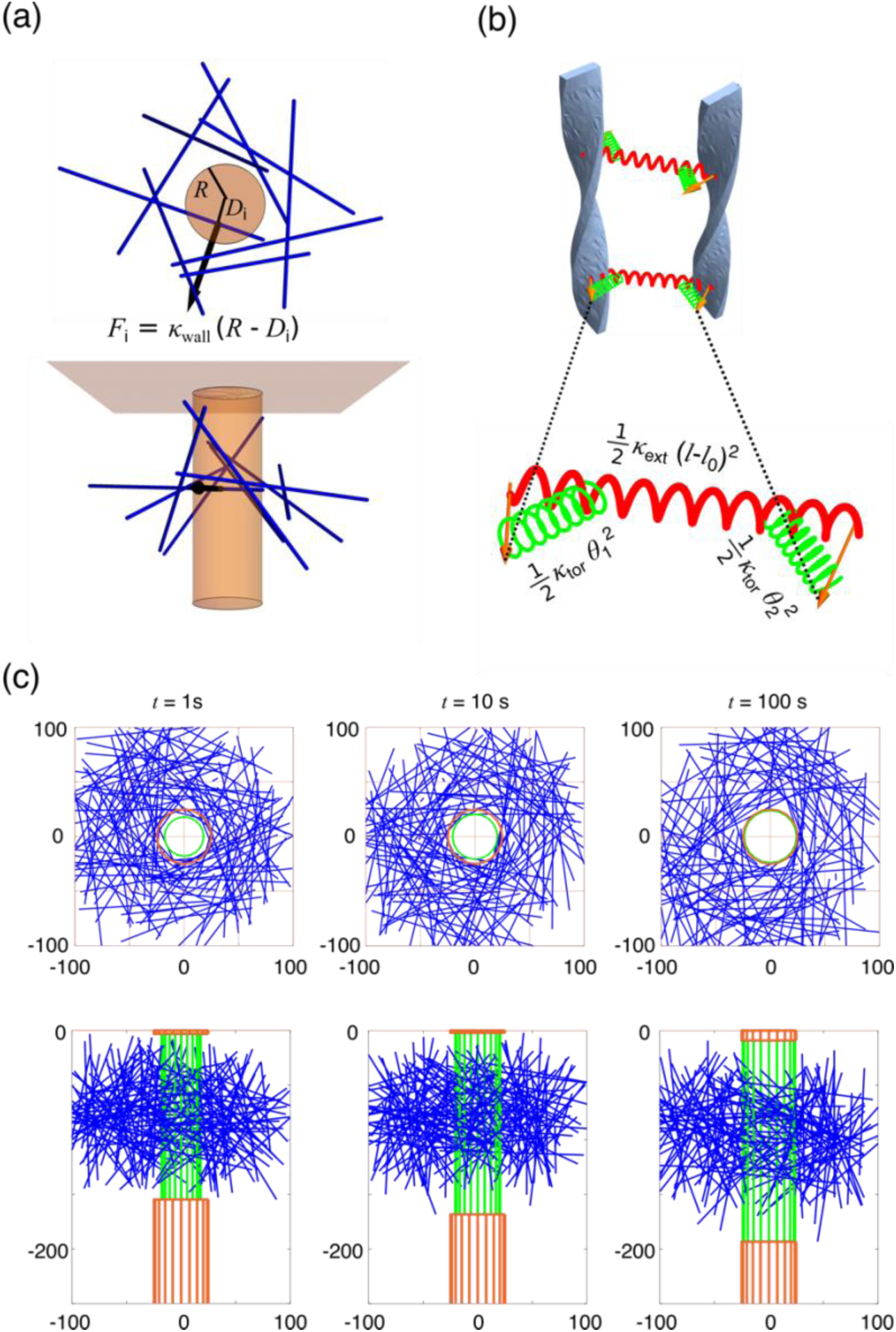
Model of a crosslinked actin network around an elastic cylinder. (a) Illustration of the model. A repulsive force is applied to filaments that penetrate into the cylinder. (b) Actin filaments are modeled as a helical rod with the arrows representing the orientation of the monomers’ binding surface with crosslinkers. Each crosslinker is modeled as a combination of one extensional spring and two torsional springs. (c) Snapshots of a typical simulation using the reference parameter values (Table 1). Each filament is depicted as a blue line. For clarity crosslinkers are not plotted. The deformed part of the cylinder (green) represents the largest cylindrical region without any filaments penetrated in.

To model actin filaments and crosslinkers, we used the same model as in Ref. (Ma and Berro 2018). In summary, each actin filament is modeled as a helical rod with a constant length *L* = 135 nm As the length of filaments is much shorter than the persistent length, we treat the filaments as a rigid body, whose movement is governed by the equation:

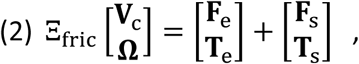

where Ξ_fric_ denotes the friction coefficient matrix, **V**_c_/**Ω** denotes the translational/rotational velocity of the filament, **F**_e_/**T**_e_ denotes the deterministic force/torque generated by the crosslinkers and the elastic cylinder, **F**_s_/ **T**_s_ denotes the stochastic force/torque due to thermal fluctuations. The helical nature of filaments is explicitly taken into account by attaching each monomer *i* of the filament with a unit vector ***m***_*i*_ representing the normal direction of the binding surface with crosslinkers (Fig. 1 b). A crosslinker is modeled as a combination of one extensional spring and two torsional springs. The extensional energy reads

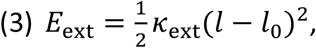

where *κ*_ext_ denotes the extensional stiffness, *l* denotes the length, and *l*_0_ denotes the rest length of the crosslinker. The torsional energy reads

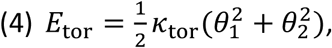

**Table 1:** List of parameters.

| Symbol | Meaning | Reference value | Varied range |
| --- | --- | --- | --- |
| $L$ | Filament length | 135nm | 135nm |
| $b$ | Filament diameter | 6nm | 6nm |
| $\delta$ | Length elongation of filaments by incorporation of a monomer | 2.7nm | 2.7nm |
| $\kappa_{\text{ext}}$ | Extensional stiffness of the crosslinker | 0.1 pN/nm | 0.1-10 pN/nm |
| $\kappa_{\text{tor}}$ | Torsional stiffness of the crosslinker | 10 pN · nm | 1-100 pN · nm |
| $R_b$ | Crosslinker binding rate constant between two unoccupied monomers | 1 s<sup>-1</sup> | 1 s<sup>-1</sup> |
| $R_u^0$ | Force-torque-free unbinding rate of a formed crosslinker | 0.1 s<sup>-1</sup> | 0.001-0.1 s<sup>-1</sup> |
| $l_0$ | Rest length of the crosslinker | 10nm | 10nm |
| $d_c$ | Reaction distance to build a crosslinker | 20nm | 20nm |
| $E_c$ | Characteristic energy of a crosslinker | 10 $k_B T$ | 10 $k_B T$ |
| $f_{\text{st}}$ | Repulsive force due to steric interaction | 100pN | 100pN |
| $k_B T$ | Thermal energy unit at room temperature | 4.1 pN · nm | 4.1 pN · nm |
| $r_{\text{min}}$ | Minimum distance between two filaments | 6nm | 6nm |
| $N_{\text{fil}}$ | Total number of actin filaments | 140 | 140 |
| $g_{\text{max}}$ | Maximum crosslinker occupancy of a filament | 0.25 | 0.25 |
| $\eta$ | Viscosity of the medium | 10<sup>-5</sup>pN · s · nm<sup>-2</sup> | 10<sup>-5</sup>pN · s · nm<sup>-2</sup> |
| $\Delta t$ | Simulation time step | 10<sup>-5</sup>s | 10<sup>-5</sup> – 10<sup>-6</sup> s |

where *κ*_tor_ denotes the torsional stiffness, *θ*_1_ and *θ*_2_ denote the angle spanned between the crosslinkers’s orientation and the attached monomer’s orientation.

A crosslinker can be formed between two monomers in two filaments with a rate constant *R*_b_ if the distance between the monomers is less than a critical value *d*_c_ and neither of the monomers is occupied. The unbinding rate of a crosslinker depends on the total elastic energy *E* = *E*_ext_ + *E*_tor_ via:

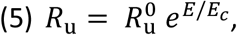

where *E*_c_ denotes the characteristic energy stored in a crosslinker. A maximum occupancy ratio *g*_max_ of crosslinkers binding to filaments is set to ensure that the total number of crosslinkers is no more than 900, which is the peak number of fimbrin during CME (Sirotkin et al. 2010).

Each simulation is run for 200s and the parameters are listed in Table 1. A more detailed description of the model can be found in Ref. (Ma and Berro 2018).

The code used to perform the simulations is available at https://github.com/ruima86/ActinAroundCylinder

## Results

### 1. Crosslinking actin filaments around a cylinder produces compressive force

We start the simulation with filaments randomly positioned and oriented around the cylinder. Crosslinks rapidly form between filaments such that a meshwrok forms around the cylinder (Fig.1 c, Movie S1). The elasticity of the crosslinkers tend to pull filaments together. However, the cylinder in the center repels filaments that penetrate into it. As a consequence, the meshwork generates a compressive force on the cylinder. The total compressive force acting on the cylinder is calculated as the sum of the magnitude of the repulsive force experienced by each filament penetrating into the cylinder. Note that the stiffness and the radius of the cylinder have a minor impact on the estimation of the forces (Figure S1). In the following, we investigate how this compressive force depends on the turnover rate and the mechanical properties of crosslinkers.

### 2. The compressive force generated by a crosslinked actin network decays over time if crosslinkers undergo turnover

If the crosslinking unbinding rate 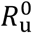 is set to zero, a positive compressive force can persist over the entire simulation, i.e. hundreds of seconds (Fig. 2a). However, when allowing crosslinkers to turnover after *t*= 10 *s* by setting their unbinding rate constant 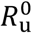 to a positive value, the compressive force decays over time (Fig. 2b and c). The decay profile has an initial rapid drop due to sudden breakage of many crosslinkers and is followed by a relatively slow decay that can be fit by an exponential function. The relaxation time of the exponential decay decreases with the unbinding rate 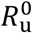.This is reflected from the fact that the dynamic force *F*_d_ at *t* = 20*s,* at which crosslinker turnover has been allowed for 10s, decreases with the unbinding rate 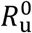, while the stable force *F*_*s*_ at *t* = 5*s* remains unchanged (Fig. 2d).

**Figure 2:**
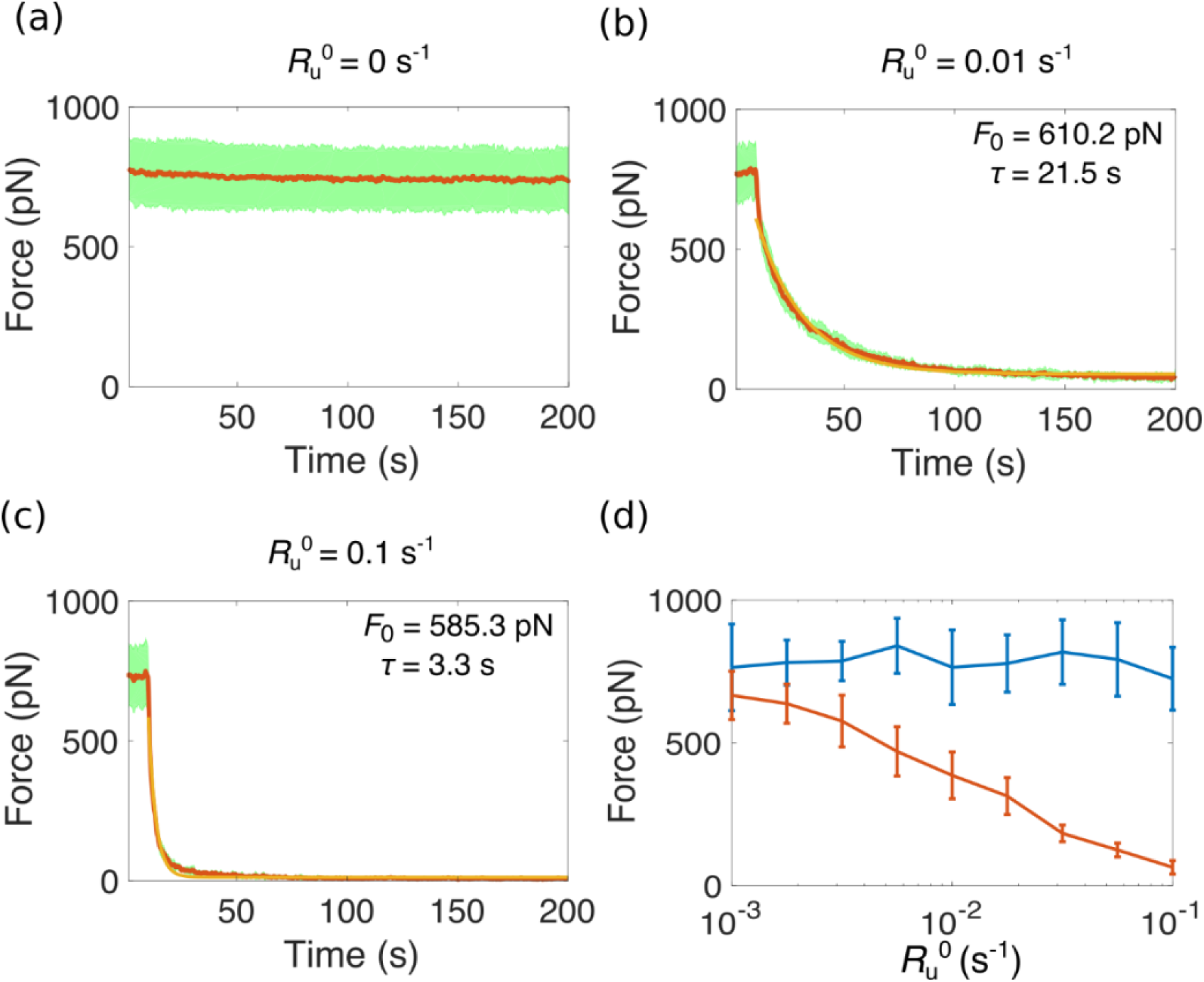
The total force decays overtime when crosslinkers are allowed toturn over. (a-c) Temporal evolution of the total force for force-torque-free unbinding rate 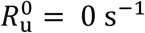 in (a), 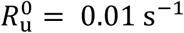 in (b) and 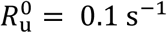 in (c). The red curve represents the average over 10 simulations and the shaded region (green) spans the standard deviation. The orange curve is the exponential fit *a e*^-(*t*-10)/*τ*^+*b* to the red curve from *t* = 10s before which crosslinker turnover is switched off such that 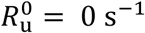 for all conditions. Inset labels show the relaxation time *τ* and the force *F*_0_ = *a* + *b* extracted from the exponential fitting. (d) Total force at *t*=5 s (blue) and *t*=20 s (red) for different unbinding rates 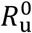.

### 3. The compressive force increases with the extensional stiffness κ_ext_ of the crosslinker

To demonstrate the compressive force originates from the elastic interaction between filaments mediated by crosslinkers, we investigate the effect of the extensional stiffness κ_ext_ of the crosslinker on the magnitude and decay time of the compressive force. Both the stable force *F*_s_ at *t*= 5s and the dynamic force *F*_d_ at *t*= 20*s* increase with κ_ext_ and can be up to 8000pN for *F*_s_ and 2000pN for *F*_d_ (Fig. 4d). The relaxation time also increases with κ_ext_. Therefore a crosslinker with high extensional stiffness is needed to produce large and persistent compressive force.

**Figure 3.**
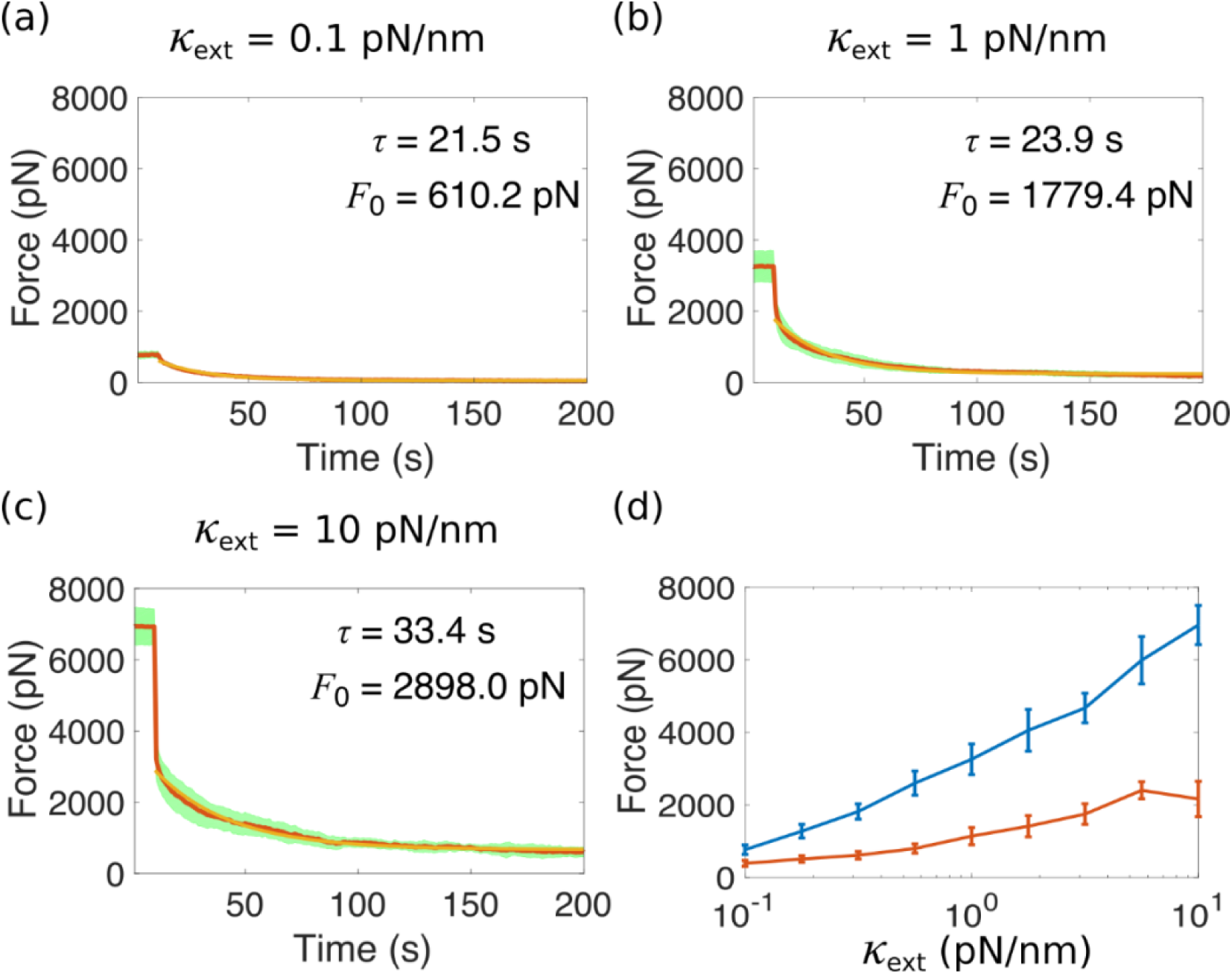
The total force generated by crosslinked actin networks strongly depends on the extensional stiffness of the crosslinker. (a-c) Temporal evolution of the total force for extensional stiffness *κ*_tor_= 1 pN·nm in (a), *κ*_tor_= 10 pN nm in (b) and *κ*_tor_= 100 pN·nm in (c). The red curve represents the average over 10 simulations and the shaded region (green) spans the standard deviation. The orange curve is the exponential fit *a e*^-(*t*-10)/*τ*^+*b* to the red curve from *t*=10s before which crosslinker turnover is switched off such that 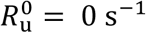 for all conditions. Inset labels show the relaxation time *τ* and the force *F*_0_ = *a* + *b* extracted from the exponential fitting. (d) The total force at *t*=5 s (blue) and *t*=20 s (red) for different extensional stiffness of the crosslinker *κ*_tor_.

**Figure 4.**
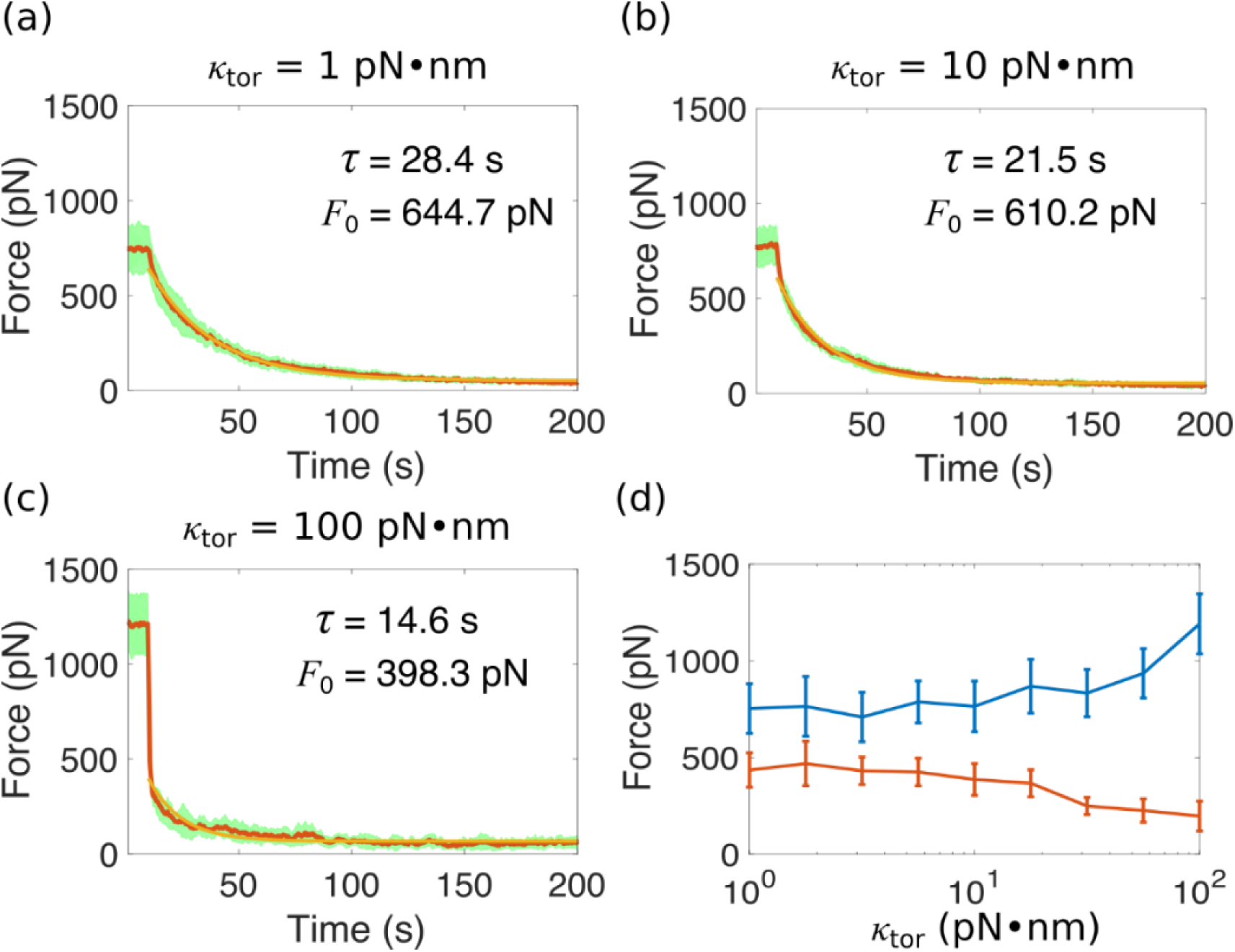
The total force generated by crosslinked actin network weakly depends on the torsional stiffness of the crosslinker. (a-c) Temporal evolution of the total force for torsional stiffness *κ*_tor_= 1 pN·nm in (a), *κ*_tor_= 10 pN·nm in (b) and *κ*_tor_= 100 pN·nm in (c). The red curve represents the average over 10 simulations and the shaded region (green) spans the standard deviation. The orange curve is the exponential fit *a e*^-(*t*-10)/*τ*^+*b* to the red curve from *t*=10s before which crosslinker turnover is switched off such that 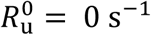 for all conditions. Inset labels show the relaxation time *τ* and the force *F*_0_=*a*+*b* extracted from the exponential fitting. (d) The total force at *t*=5 s (blue) and *t*=20 s (red) for different torsional stiffness of the crosslinker *κ*_tor_.

### 4. The torsional stiffness κ_tor_ of the crosslinker has a weak effect on the compressive force

With increasing torsional stiffness κ_tor_ of the crosslinker, both the stable force *F*_s_ and the dynamic force *F*_d_ remain almost flat until κ_tor_ = 1.8 · pN·nm. The stable force *F*_*s*_ then increases with κ_tor_ but the dynamic force *F*_d_ shows opposite trend. This is likely due to the fact that with increasing κ_tor_, the torsional energy stored in crosslinkers increases. Once crosslinker turnover is switched on, the stored elastic energy is rapidly released to relax the stress and a higher κ_tor_ leads to a higher effective unbinding rate of the crosslinkers, since it increases exponentially with the stored energy (Equation 5).

## Discussion

In this paper, we have studied the compressive force produced by crosslinked actin filaments. We showed that a crosslinked actin network generates force if the filaments are placed around an elastic cylinder. The force is compressive and decays over time as a result of crosslinker turnover. The magnitude and persistent time of the force strongly depends on the extensional stiffness κ_ext_ of the crosslinkers but weakly on their torsional stiffness κ_tor_. More than a thousand piconewtons of force can be produced by crosslinkers with an extensional stiffness κ_ext_ = 10 pN/Nm and last for tens of seconds. Since formation of a clathrin-coated pit is fast (∼10 s), this force could be a significant contribution to the force required for endocytosis, especially when compared with the force that can be produced by the polymerization of the same number of actin filaments (∼150 pN) (Kaksonen, Toret, and Drubin 2006; Lacy et al. 2018). However, the force we have shown here is a compressive force, which plays a role of line tension but does not contribute to the directed force needed to pull a membrane inward. This line tension might help deform the membrane through a buckling mechanism (Lenz, Crow, and Joanny 2009). A rough estimation by dimensional analysis of the buckling line tension *F*_buckle_ for a flat membrane patch with a base radius of *r*= 50nm (Kukulski et al. 2012) and bending rigidity of κ= 2000 *k*_B_*T* (Dmitrieff and Nédélec 2015), and ignoring the contribution of turgor pressure, leads to 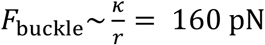. Thereforethe compressive force generated by the crosslinked actin network is around an order of magnitude larger than the force required to buckle a flat membrane.

In this study, we have neglected actin dynamics but focused on the force production mechanism by passively crosslinking actin filaments. The energy source comes from imposing an initially highly crosslinked actin network. This is a state which is out of equilibrium and the decay of the force is essentially the relaxation process for the system to reach an equilibrium. When actin dynamics is considered, the system will be driven out of equilibrium, and we expect that the actin meshwork will be able to sustain force production for a longer time.

## Acknowledgements

The authors have no conflict of interest to declare. This research was supported by the National Institute of Health/National Institute of General Medical Sciences Grant R01GM115636.

## Supplemental Figure

**Figure S1:**
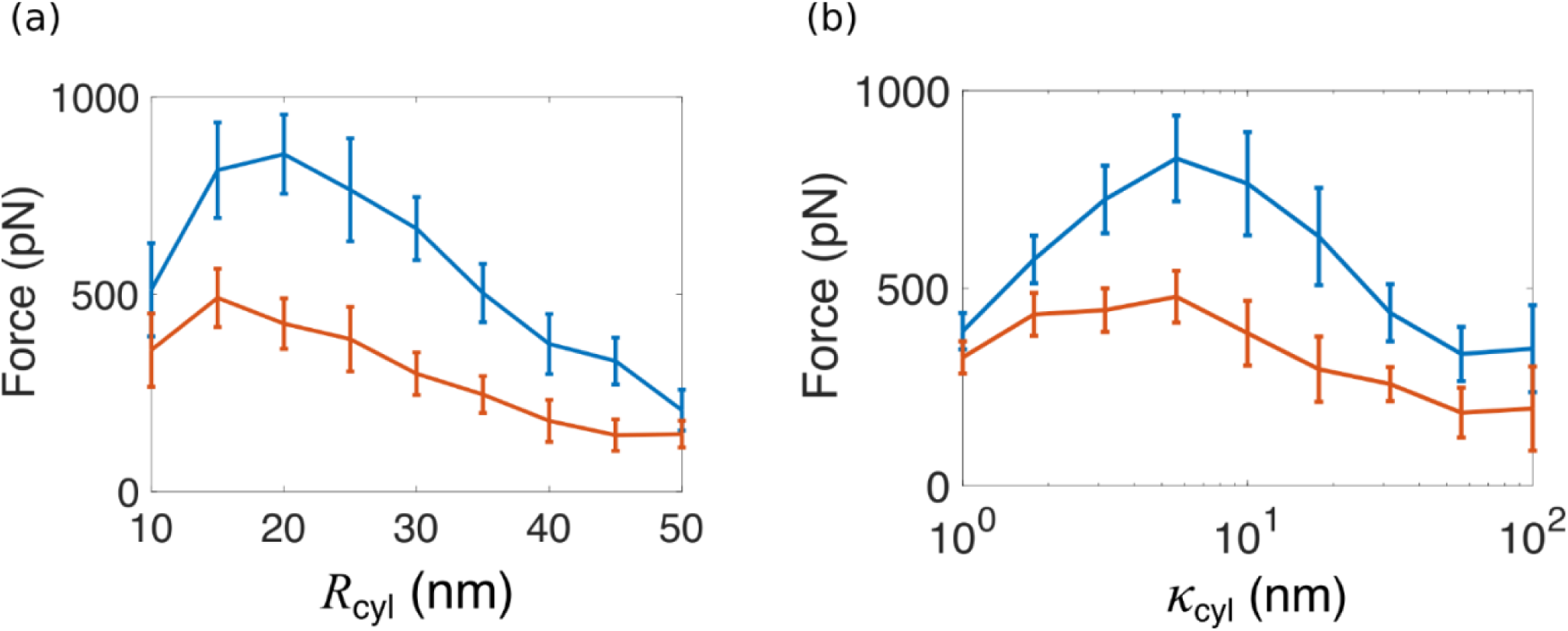
Effect of the radius and the stiffness of the cylinder on the compressive force. (a) The total force at *t*=5 s (blue) and *t*=20 s (red) for different radii of the cylinder *R*_cyl_. (b) The total force at *t*=5 s (blue) and *t*=20 s (red) for different stiffnesses of the cylinder *R*_cyl_.

## Supplemental movie

**Movie S1: Simulation of actin filaments around an elastic cylinder.** (a-b) Each filament is represented by a blue line. For clarity, crosslinkers are not plotted. The cylinder is divided in two parts, the green part represents the region where actin filaments penetrate inside the cylinder, the orange part is void of filaments. The radius of the green part represents the minimum distance of all the filaments penetrated into the cylinder to the central axis of the cylinder. The simulation is run for 200s. (a) Side view. (b) Top view.

