## Supplementary figures and images for "Crosslinking actin networks produces compressive force"

### Supplemental figure S1

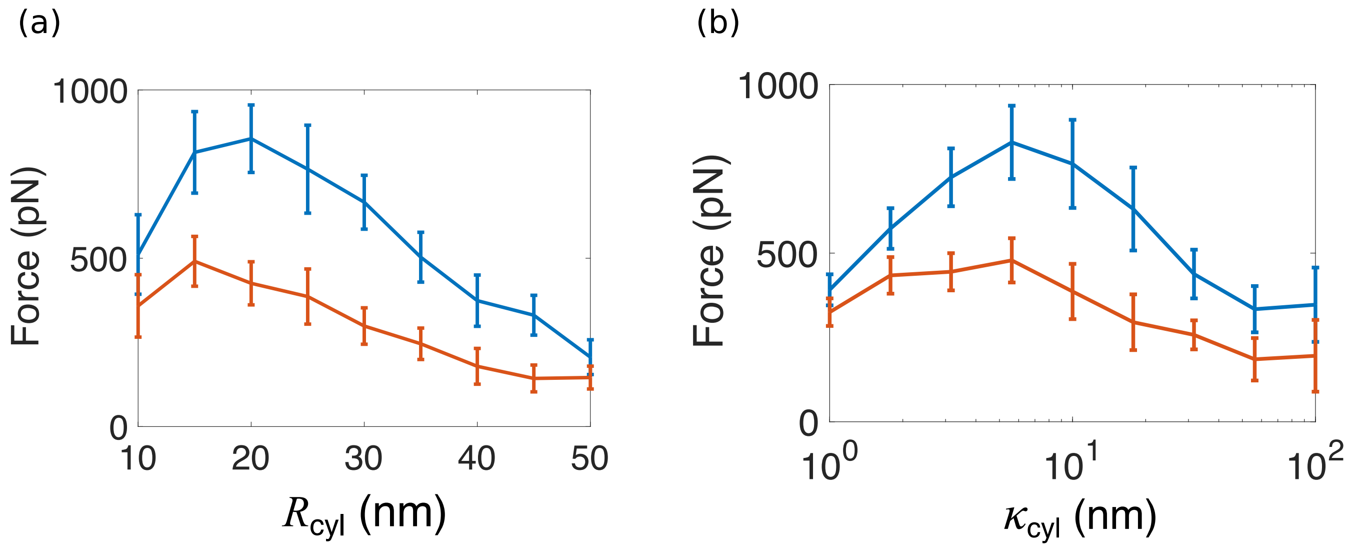
